# Human progenitor T-cell differentiation regulated by the mechanical resistance of thymus-mimetic extracellular matrices

**DOI:** 10.1101/2025.08.27.671384

**Authors:** Nicholas Jeffreys, Kyle T. Ruark, Joshua M. Price, Ella M. Serrano-Wu, Blake Hanan, Andrew S. Khalil, Wei-Hung Jung, Nuria Lafuente-Gómez, Joshua M. Brockman, Andrew Lu, Izabela Zmirska, Kyle H. Vining, Junzhe Lou, Kwasi Adu-Berchie, Siyoon Kwon, Hamza Ijaz, Azeem Sharda, David T. Scadden, David J. Mooney

## Abstract

Therapeutic T-cell engineering *ex vivo* from human hematopoietic stem cells (HSCs) focuses on recapitulating notch1-signaling and α4β1-integrin-mediated adhesion within the thymic niche with supportive stromal cell feeder-layers or surface-immobilized recombinant protein-based engineered thymic niches (ETNs). The relevant Notch1-DLL-4 and α4β1-integrin-VCAM-1 interactions are known to respond to mechanical forces that regulate their bond dissociation behaviors and downstream signal transduction, yet manipulating the mechanosensitive features of these key receptor-ligand interactions in thymopoiesis has been largely ignored in current ETN designs. Here, we demonstrate that human T-cell development from cord blood-derived CD34^+^ HSCs is regulated via molecular cooperativity in notch1 and integrin-mediated mechanotransduction. Mechanically confining interpenetrating network (IPN) hydrogel-based 3D cell culture comprised of collagen type I and alginate polysaccharides functionalized with DLL-4 and VCAM-1 is used as a model viscoelastic 3D ETN to manipulate human progenitor (pro)T-cell differentiation. This ETN enables orthogonal control of the mechanical and biomolecular features of the thymic niche, including thymopoietic ligand density, modulus, and viscoelastic properties (e.g., stress relaxation kinetics). We identify that soft, viscous matrices that enhance activation of the notch1-pathway, and subsequently notch1 intracellular domain (NICD) nuclear import sustain the T-cell development gene regulatory network during proT-cell differentiation. Conversely, stiff, elastic matrices inhibit HSC commitment to the T-lineage, and rather promotes Myeloid-cell differentiation. Our observations indicate mechanical reciprocity in signaling pathways indispensable to thymopoiesis, and highlights extracellular matrix mechanics as a variable in controlling hematopoietic stem cell fate decisions.

## Introduction

T-cell engineering *ex vivo* from human hematopoietic stem cells (HSCs) focuses on recapitulating indispensable features of thymopoiesis (e.g. notch1-signaling^1^ and α4β1-integrin-mediated adhesion^2^) through either supportive stromal cell feeder-layers that ectopically express the notch1 ligand DLL-4 or 2D surface immobilized DLL-4 and VCAM-1^3–5^. Synergy between the notch1-pathway and integrin-mediated adhesion via DLL-4 and VCAM-1 has been reported to enhance human progenitor (pro)T-cell differentiation velocity, but how integrin-substrate adhesions support notch1-signaling in the context of thymopoiesis remains elusive^6^. Additionally, developing thymocytes interact with thymic epithelial cells within a mechanically confining nanoporous extracellular matrix (ECM)^7–9^ that exhibits viscoelastic properties such as stress relaxation behavior^10,11^, but knowledge of the combinatorial effects of ECM mechanics (e.g., modulus, stress relaxation behavior, and the mechanical coupling of thymocytes to the ECM via integrin-ECM adhesions) on thymopoiesis is incomplete^12–15^.

Notch1-DLL-4 interactions and integrin-substrate adhesions are anisotropic mechanosensors that respond to piconewton (pN)-level forces that tune ligand sensitivity and receptor-ligand mechanochemical equilibria^16–20^. Specifically, tensile load applied to the notch1-ligand bond permits proteolytic cleavage and release of the notch1 intracellular domain (NICD) transcription factor into the cytosol to translocate to the nucleus and act on downstream target genes^17,21^. Integrin-ligand bonds respond to the mechanical properties of their substrate to mediate cellular signals such as cell fate decisions^22–24^, cytoskeletal organization^25,26^, and the nuclear import of mechanosensitive transcription factors^27–29^. Interestingly, substrate mechanics in concert with substrate-tethered notch1 ligands has been reported to impact cellular behavior and the differential expression of downstream notch1 target genes^12,30,31^. Despite these observations, the biomolecular mechanisms underlying mechanical regulation of the notch1 pathway by substrate mechanics and its cooperation with actomyosin contractility via integrin-substrate adhesions remain poorly understood.

Here, we hypothesized that substrate mechanics may regulate human T-cell development from cord blood-derived CD34^+^ HSCs. Mechanically confining interpenetrating network (IPN) hydrogel-based 3D cell culture comprised of collagen type I and alginate polysaccharides functionalized with DLL-4 and VCAM-1 is used as a model viscoelastic 3D engineered thymic niche (ETN) to manipulate human progenitor (pro)T-cell differentiation. This ETN permits decoupling of the mechanical and biomolecular features of the thymic niche, including thymopoietic ligand density, modulus, and viscoelastic properties (e.g., stress relaxation kinetics). Through combinatorial small molecule inhibition of the notch1 pathway and actomyosin contraction, we identify that soft, viscous matrices enhance human proT-cell differentiation velocity. Interestingly, stiff, elastic matrices inhibit HSC commitment to the T-lineage, and rather promotes Myeloid-cell differentiation. These observations indicate mechanical cooperation between notch1-signaling and integrin-substrate adhesions within the context of thymopoiesis, and highlights extracellular matrix mechanics as a variable in controlling hematopoietic stem cell fate decisions within their cellular niche. These findings may have utility for future *in vitro* models of thymopoiesis and therapeutic T-cell biomanufacturing strategies from stem cells.

## Results

### Thymus-mimetic ECMs decouple modulus from stress relaxation behavior and ligand density

To delineate the impacts of substrate mechanics on human proT-cell differentiation, we developed a 3D semisynthetic thymus-mimetic ECM (3D ETN) in which modulus can be decoupled from viscoelastic properties (e.g., stress relaxation kinetics and plasticity) and thymopoietic ligand density (**Fig. 1a**). 3D ETNs were synthesized from interpenetrating polymer networks (IPNs) of tetrazine (Tz)-modified alginates and collagen I fibers to investigate whether encapsulated cord blood-derived CD34^+^ human hematopoietic stem cells (HSCs) responded to orthogonal permutations in substrate modulus and stress relaxation behavior^32–34^. Alginate polysaccharides are bioinert to cells^35,36^ and dictates the bulk modulus through the chelation of divalent calcium ions, while collagen I is a key matrix constituent of the thymus^7,8,12^. Alginate and collagen I concentrations were fixed in order to delineate solely the impacts of permutations in bulk modulus and stress relaxation behavior on human proT-cell differentiation, such that potential effects of collagen I ligand density and soluble protein transport through changes in matrix mesh size are ruled out (**Supplementary Table 1**)^23,29^. The degree of ionic crosslinking in the alginate matrix controls the bulk modulus of the 3D ETN (**Fig. 1c**), independent of the stress relaxation behavior which can be changed by reinforcing the alginate matrix with additional covalent crosslinks through the introduction of tetra-functional PEG crosslinkers terminated with Norbornene ‘click’ moieties (Nb) (**Fig. 1a, 1b**). Shear stress relaxation tests confirmed that the relaxation half-time of covalently crosslinked 3D ETNs was higher than that of purely ionically crosslinked 3D ETNs (**Fig. 1b, 1d**). In line with the changes in relaxation half-time, the loss angle of covalently crosslinked 3D ETNs was lower than that of purely ionically crosslinked 3D ETNs (**Fig. 1e, 1f**). Creep measurements confirmed these changes in both viscoelastic and viscoplastic properties, where purely ionically crosslinked 3D ETNs retained greater irreversible plastic strain than covalently crosslinked 3D ETNs (**Fig. 1g, 1h**). The more ‘liquid’-like 3D ETNs that exhibit faster stress relaxation kinetics are termed ‘viscous’, and the more ‘solid’-like 3D ETNS that exhibit slower stress relaxation kinetics are termed ‘elastic’ (**Fig. 1a**).

**Fig. 1.**
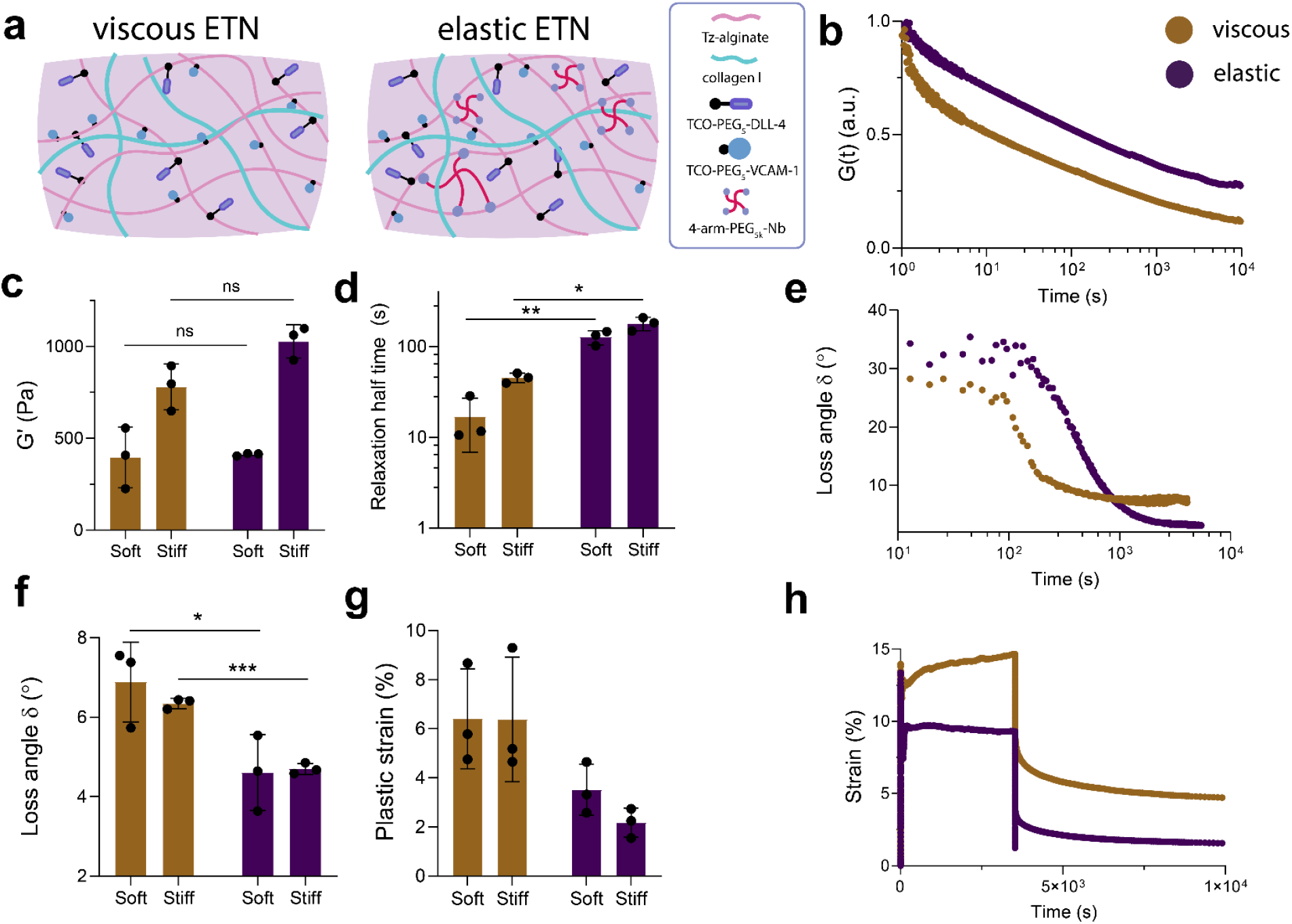
Interpenetrating network collagen I-tetrazine-alginate hydrogels functionalized with DLL-4 and VCAM-1 enable orthogonal control over the viscoelastic properties of a 3D engineered thymic niche. a) Schematic of physicochemical composition of viscous and elastic 3D ETNs. b) Stress relaxation kinetics of viscous and elastic 3D ETNs at 15% fixed shear strain. c) Storage modulus values for viscous and elastic 3D ETNs (n = 3); ns: no significance, Unpaired t test with Welch’s correction. d) Characteristic time for 3D ETN matrices to relax 50% of the initial stress in response to fixed deformation (n = 3); *p < 0.05, **p < 0.01, Unpaired *t* test with Welch’s correction and 95% confidence interval. e) Loss angle oscillatory time sweep kinetics for viscous versus elastic 3D ETN matrices. f) Loss angle measurements for 3D ETN matrices (n = 3); *p < 0.05, ***p < 0.001, Unpaired *t* test with Welch’s correction and 95% confidence interval. g) Irreversible plastic strain measurements for 3D ETN matrices (n = 3). h) Creep behavior for viscous and elastic 3D ETNs. Data is visualized as mean values +/-SD.

Trans-cyclooctene (TCO)-modified thymopoietic ligands DLL-4 and VCAM-1 were directly ligated to the Tz-alginate backbone via an inverse-electron-demand Diels-Alder cycloaddition ‘click’ protein-polymer conjugate reaction to enable rapid and highly efficient ligand conjugation (**Fig. 1a**)^37,38^. The ligand concentrations of DLL-4 and VCAM-1 were kept constant in these studies with a stochiometric ratio of 5:1 DLL-4:VCAM-1, which has been previously demonstrated to promote human proT-cell differentiation (**Supplementary Table 1**)^4,6,12^. Importantly, these fixed concentrations enable direct inference of how mechanical cues conveyed from the substrate impact notch1-DLL-4, integrin-VCAM-1/collagen I interactions, and downstream mechanotransduction in encapsulated CD34^+^ HSCs within the 3D ETN.

### Thymic-ECM mechanics differentially impacts HSC lineage commitment

The impact of 3D ETN substrate mechanics on HSC lineage fate decisions was next investigated. CD34^+^ HSCs were isolated from umbilical cord blood units procured from the Dana Farber Cancer Institute Pasquarello tissue bank and expanded in a serum-free chemically defined HSC expansion media (**Fig. 2a, Supplementary Table 5**)^41^. The differentiation potential of isolated CD34^+^ HSCs was confirmed by performing a colony forming unit assay (**Supplementary Fig. 1**). CD34^+^ HSCs were capable of forming CFU-Es, BFU-Es, CFU-GMs, CFU-GEMMs, and CFU-Ms.

**Fig. 2.**
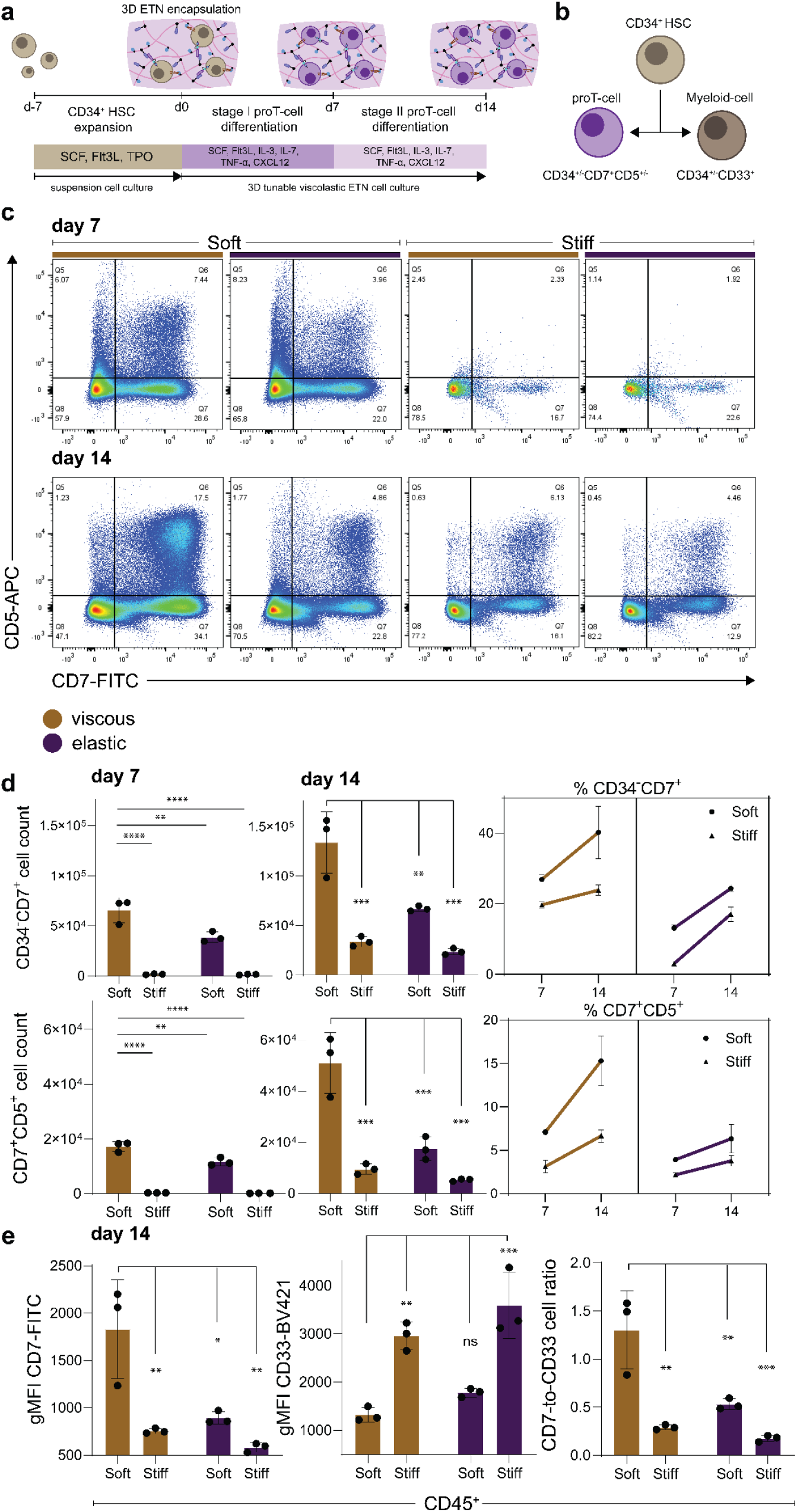
Softer, viscous 3D engineered thymic niche matrices enhance progenitor T-cell differentiation whereas stiffer, elastic matrices promote Myeloid-cell differentiation. a) proT-cell differentiation protocol in 3D ETN matrices of tunable physical properties. CD34^+^ cells were expanded (d-7-d0), encapsulated in matrices of varying viscoelasticity and stiffness (d0), and cultured in a chemically defined differentiation media (stage I media, d0-d7, stage II media, d7-d14). b) Scheme of lineage fate decisions of CD34^+^ HSCs into either CD7^+^ proT-cells or CD33^+^ Myeloid cells. c) Representative flow cytometry plots of CD7^+^CD5^+^ proT-cells as a function of 3D ETN matrix mechanics (d7 and d14). d) Cell counts and frequencies of CD34^-^CD7^+^ and CD7^+^CD5^+^ proT-cells as a function of matrix mechanics (d7 and d14) (n = 3); Two-way ANOVA with Tukey’s post-hoc test and 95% confidence interval. e) geometric mean fluorescent intensity (gMFI) of CD7 and CD33 on CD45^+^ cells and CD7-to-CD33 cell ratio as a function of matrix mechanics (d14) (n = 3); Two-way ANOVA with Tukey’s post-hoc test and 95% confidence interval. Data is visualized as mean values +/- SD. *p < 0.05, **p < 0.01, ***p < 0.001, ****p < 0.0001, ns (no significance).

Expanded CD34^+^ HSCs were encapsulated in 3D ETNs of varying viscoelastic properties (soft-viscous, soft-elastic, stiff-viscous, and stiff-elastic) and cultured in serum-free, staged, human proT-cell differentiation media (stage I (d0-d7) and stage II (d7-d14)) (**Fig. 2a, Supplementary Table 5**). Encapsulated cells were subsequently phenotyped for human proT-cell markers and Myeloid-lineage markers by flow cytometry on day 7 and day 14 of culture to assess lineage fate decisions (**Fig. 2b**). Softer and more viscous 3D ETN matrices were observed to enhance the emergence of CD7^+^CD5^+^ proT-cells compared to stiffer, more elastic matrices (**Fig. 2c**). Stiffer and more elastic matrices abrogated the differentiation kinetics of CD34^+^ HSCs towards the T-lineage, whereas softer and more viscous matrices enhanced proT-cell differentiation velocity, increasing both the gross cell count and frequency of CD7^+^CD5^+^ proT-cells (**Fig. 2d**). Cells cultured in soft-viscous 3D ETNs additionally had significantly higher expression of CD7 compared to stiffer and more elastic matrices, as indicated by the geometric mean fluorescent intensity (gMFI) (**Fig. 2e**). Interestingly, despite having exogenous thymopoietic cues presented from both the matrix and the soluble media, CD34^+^ HSCs cultured in stiffer and more elastic matrices differentiated towards the myeloid lineage (**Fig. 2e**). Analysis of CD33 gMFI and the ratio of CD7^+^ lymphoid cells to CD33^+^ myeloid cells further illustrated that CD34^+^ HSCs were significantly biased towards generating multipotent myeloid-committed progenitor cells when cultured in stiffer, elastic 3D ETNs (**Fig. 2e**).

### Notch1-signaling and actomyosin contractility synergize to enhance T-lineage commitment in soft-viscous thymus-mimetic ECMs

We next investigated whether the 3D ETNs are permissive of activating the notch1-signaling pathway at the gene expression level. Soluble notch1 ligands are incapable of activating the notch1 mechanotransduction module, as the receptor requires tensile loads in order to proteolytically release the NICD^4,39,40^. Expanded CD34^+^ HSCs were again encapsulated within ‘soft-viscous’ 3D ETNs and cultured in a serum-free, staged, human proT-cell differentiation media (**Fig. 3a, Supplementary Table 5**) to investigate if 3D ETNs are capable of initiating notch1-signaling and activating downstream notch1 target genes comprising the human T-cell development gene regulatory network (GRN) (**Fig. 3b**)^42,43^. Examination of key genes comprising the T-cell development GRN within cells after 96 hours of culture within ‘soft-viscous’ 3D ETNs compared to pre-encapsulated CD34^+^ HSCs revealed upregulation of downstream targets of NICD (TCF7, Hes1, Deltex, Notch1) (**Fig. 3c, 3d**)^44^. In contrast, the myeloid-lineage transcription factor Pu.1 was downregulated, along with the HSC self-renewal gene E2a (**Fig. 3c, 3d**)^45,46^. Therefore, 3D ETNs permit activation of the notch1 pathway to initiate thymopoiesis through regulation of the T-cell development GRN.

**Fig. 3.**
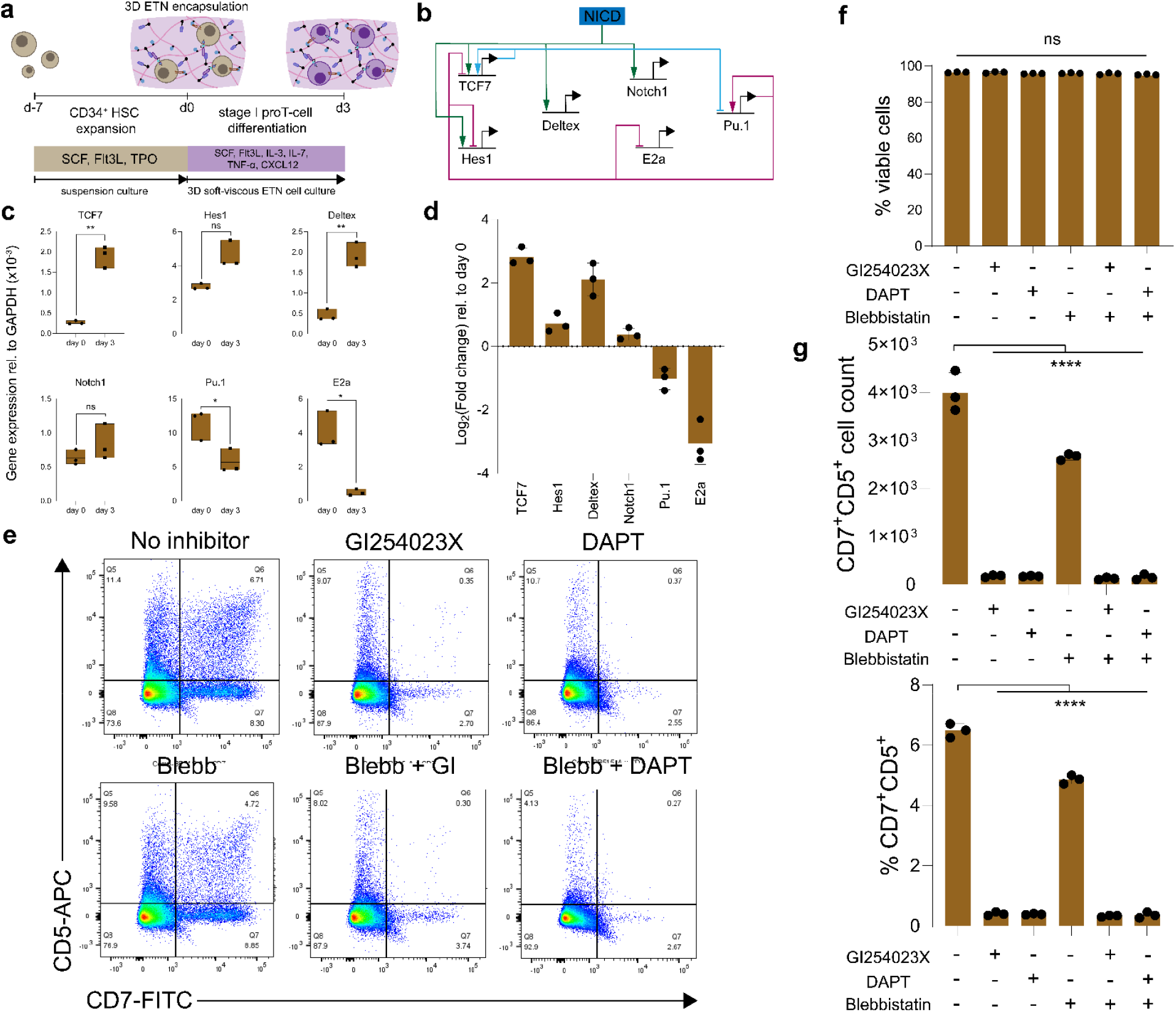
Notch1-signaling and actomyosin contractility synergize to promote progenitor T-cell differentiation. a) Total RNA was isolated from expanded CD34^+^ HSCs on day 0 prior to 3D ETN encapsulation and from encapsulated cells in soft-viscous ETN matrices cultured in stage I proT-cell differentiation media on day 3. b) NICD shuttles to the nucleus to sustain the T-cell development gene regulatory network comprising several feedback network motifs. c) Gene expression of representative notch1 target genes (TCF7, Hes1, Deltex, Notch1), a Myeloid-lineage commitment gene (Pu.1), and an HSC self-renewal gene (E2a) (n = 3); *p < 0.05, **p < 0.01, ns (no significance), Unpaired *t* test with Welch’s correction and 95% confidence interval. d) Log_2_(Fold change) in gene expression in cells that were encapsulated in soft-viscous matrices for 3 days relative to expanded CD34^+^ HSCs on day 0 (n = 3). e) Representative flow cytometry plots of CD7^+^CD5^+^ proT-cells as a function of combinatorial small molecule inhibitor treatment (d7). f) % cell viability as a function of combinatorial small molecule inhibitor treatments (d7) (n = 3); One-way ANOVA with Dunnett’s post-hoc test and 95% confidence interval. g) Cell counts and frequencies of CD7^+^CD5^+^ proT-cells as a function of combinatorial small molecule inhibitor treatment (d7) (n = 3); One-way ANOVA with Dunnett’s post-hoc test (compared to No inhibitor treatment, 95% confidence interval. Data is visualized as mean values +/- SD; ****p < 0.0001, ns (no significance).

As notch1-signaling and integrin-substrate adhesions have been demonstrated to synergize to enhance proT-cell differentiation velocity, we sought to investigate the underlying biomolecular mechanisms underlying notch1/integrin mechanical cooperativity. Previous studies have suggested that actomyosin contractility may support notch1-signaling^47,48^. Actomyosin has extensive lateral contacts with the adherent substrate across the cell membrane through integrin-substrate coupling, and is postulated to enable receptor-mediated endocytosis and myosin-dependent tension to support the tensile load-dependent proteolytic activation of the notch1 pathway^17,21,47^. Additionally, the presence of actomyosin at the nuclear periphery and its co-localization with the linker of nucleoskeleton and cytoskeleton (LINC) complex protein Nesprin2 and apical actin caps suggests that actomyosin contraction may support the nucleocytoplasmic transport of NICD and other macromolecular cargoes important in sustaining the T-cell development GRN by increasing nuclear influx through deformation of the nuclear pore complex (NPC)^28,47,49,50^.

We therefore performed a combinatorial small molecule inhibition study targeting actomyosin contractility and critical molecular features of notch1-signaling on CD34^+^ HSCs encapsulated in ‘soft-viscous’ 3D ETN matrices (**Supplementary Table 3**). Selective inhibitors targeting ADAM10-mediated notch1 cleavage (GI254023X (GI))^51^ and γ-secretase-mediated NICD release (DAPT)^42^ were used in concert with an inhibitor targeting myosin II contractility (Blebbistatin (Blebb)) in order to delineate potential synergies between notch1 and integrin-mediated mechanotransduction. An inhibitor-dose response study on suspension culture CD34^+^ HSCs determined maximal working concentrations for each inhibitor (**Supplementary Fig. 3, Supplementary Table 3**). Consistent with previous reports, GI and DAPT selectively inhibited proteolytic cleavage of the notch1 receptor while binding to DLL-4, significantly obstructing the emergence of CD7^+^CD5^+^ proT-cells (**Fig. 3e**). Flow cytometry analysis revealed minimal cytotoxicity (≥ 95% cell viability) of small molecule inhibitor combinations on cells encapsulated in soft-viscous 3D ETNs (**Fig. 3f**). Blebb decreased CD7^+^CD5^+^ proT-cell emergence, whereas combinations of Blebb and GI or DAPT significantly obstructed CD7^+^CD5^+^ proT-cell emergence (**Fig. 3e**). Analysis of the gross cell count and frequency of CD7^+^CD5^+^ proT-cells after 7 days of culture in soft-viscous 3D ETNs further supported this observation, demonstrating significant knockdown of CD7^+^CD5^+^ proT-cell emergence when cells were treated with GI, DAPT, Blebb, GI + Blebb, and DAPT + Blebb (**Fig. 3g**).

## Discussion

Altogether, these experimental findings suggest that CD34^+^ HSC fate decisions within the thymic niche can be controlled through the mechanical manipulation of cell-matrix interface in a 3D ETN that physically presents DLL-4, and integrin binding motifs (VCAM-1 and collagen I) from the substrate. Soft-viscous matrices enhanced the emergence of CD7^+^CD5^+^ proT-cells from their hematopoietic precursors, whereas stiffer and more elastic matrices promoted HSC lineage commitment towards the myeloid lineage (**Fig. 2b-2e**). The impact of stiff, elastic substrates in the promotion of Myeloid cell generation and their downstream progeny has been previously established, though the underlying mechanisms driving HSC lineage fate decisions towards the myeloid-fate as a function of ECM mechanics still remains elusive^33,55,56^.

These studies also suggest that synergies between notch1-signaling and actomyosin contractility (via integrin-VCAM-1/collagen I adhesions) enhance proT-cell differentiation velocity. Notch1-DLL-4 interactions and integrin-substrate adhesions implicate a ‘catch’ bond that tunes ligand sensitivity, where optimal tensile loads applied to the bond sustains the bond lifetime to enhance the probability of notch1 cleavage by ADAM10 and γ-secretase to release NICD into the cytosol, and the formation of integrin focal adhesion complexes in response to substrate mechanics^19–21,57^. Soft-viscous matrices may permit greater notch1-DLL-4 and integrin-substrate bond formation, and enable the energization of the notch1-DLL-4 catch bond through passive injection of energy by actomyosin contraction through integrin substrate bonds and perturbations in thymocyte membrane dynamics (**Fig. 4**)^17,47^. Stiffer and more elastic matrices could promote load-induced slippage, leading to rapid dissociation of notch1-DLL-4 bonds and focal adhesion complexes. In the absence of notch1-signaling, encapsulated CD34^+^ HSCs preferentially differentiate into myeloid-competent cells through asymmetric division^58,59^. Moreover, in concert with notch1-signaling, actomyosin contraction may also enable cytokinesis and asymmetric division of HSC progenitors into proliferating thymocytes that plastically deform the soft, viscous matrix in a feed forward loop to enhance thymocyte maturation (**Fig. 3e-3g, Fig. 4**)^60–63^. Combinatorial small molecule inhibition of notch1 cleavage and actomyosin contraction within cells encapsulated in soft-viscous 3D ETNs also inhibited generation of CD7^+^CD5^+^ proT-cells (**Fig. 3g**).

**Fig. 4.**
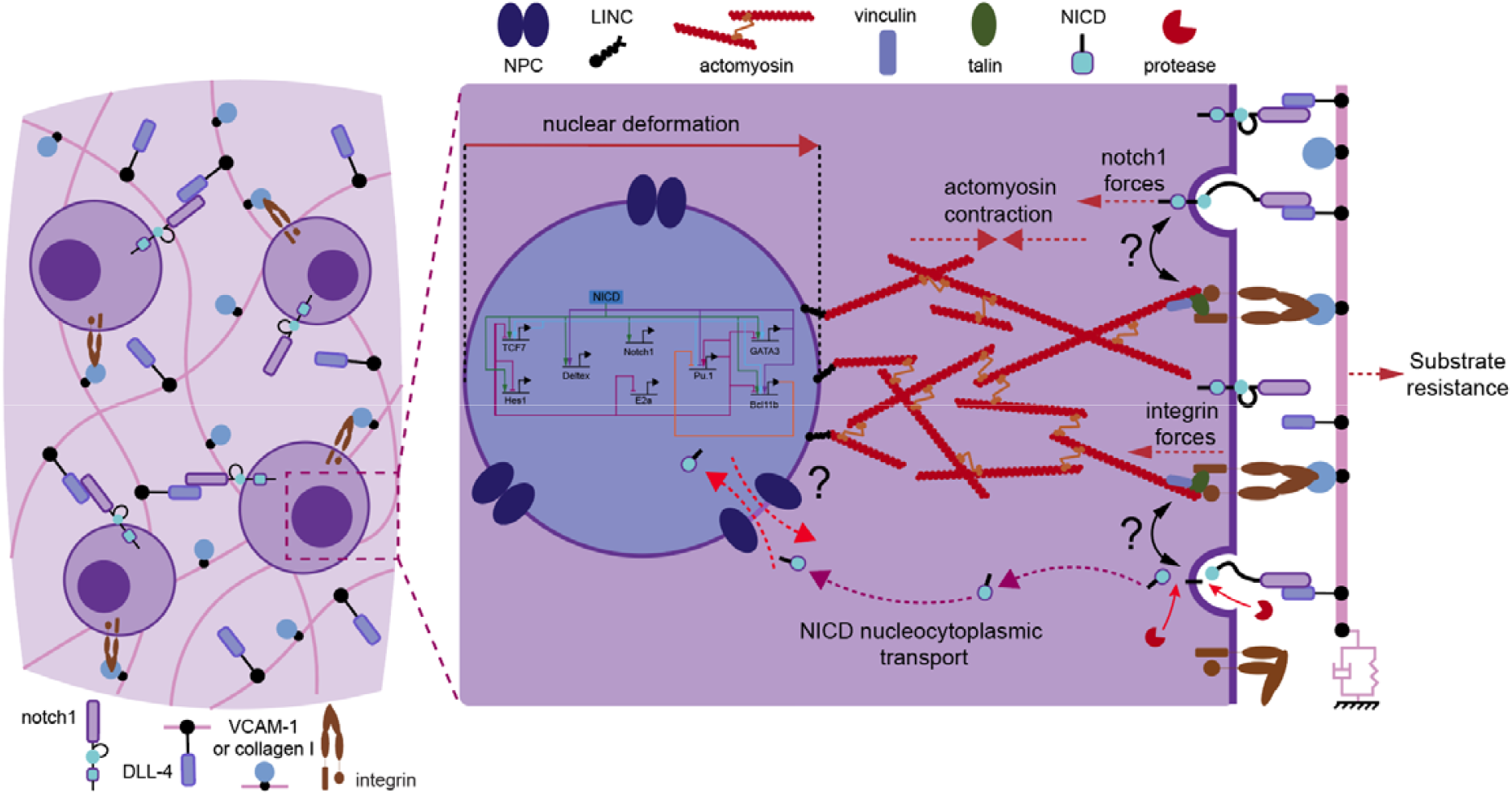
Potential mechanical determinants of Notch1-signaling and actomyosin contractility synergy in promoting progenitor T-cell differentiation. At the cell-substrate interface, softer, viscous matrices promote greater ‘catch’ bond engagement of notch1 to DLL-4 and integrins to VCAM-1 and collagen I, increasing the probability of proteolytic notch1 cleavage to release NICD into the cytosol. Actomyosin contraction via integrin-substrate adhesions directly energizes the notch1-DLL-4 catch bond through perturbations in membrane dynamics and cortical actomyosin tension, and enables tensile loading upon notch1 through receptor-mediated endocytosis. This deforms the nonregulatory region of the notch1 receptor to enable proteolytic cleavage by ADAM10 and γ-secretase. In concert with notch1-signaling, actomyosin contraction enables cytokinesis and asymmetric division of HSC progenitors into proliferating thymocytes that plastically deform the soft, viscous matrix in a feed forward loop to enhance thymocyte maturation. Actomyosin-nucleoskeletal coupling at the nucleus may potentially increase the flux of NICD across the NPC to sustain the T-cell development GRN in a feed forward loop that promotes thymocyte maturation. NICD: notch1 intracellular domain, LINC: nucleoskeleton and cytoskeleton, NPC: nuclear pore complex.

Collectively, these observations emphasize potential mechanical determinants of signaling pathways indispensable to thymopoiesis, and highlights ECM mechanics as a variable in controlling hematopoietic stem cell fate decisions within their cellular niche. These findings may therefore have additional utility for future *in vitro* models and investigations of thymopoiesis, and for T-cell biomanufacturing strategies from stem cells.

## Materials and Methods

### Materials synthesis, preparation, and storage

Low-molecular-weight (LMW, M_W_) ultrapure very-low-viscosity-G block enriched (VLVG) sodium alginate (Pronova UP VLVG alginate, NovaMatrix, BP-1802-03) was used as a base polymer for these studies, with an approximate M_W_ of 32kDa, a glucuronic acid to mannuronic acid (G/M) ratio of >1.5, and <100 endotoxin units per gram of alginate, as according to manufacturer specifications.

Tetrazine-(Tz)modified UP VLVG alginate was synthesized by covalent coupling of (4-(6-methyl-1,2,4,5-tetrazin-3-yl)phenyl)methanamine hydrochloric acid (Tz-HCl, KareBay Biochem, Inc., KB220830) through aqueous carbodiimide chemistry, as previously described to yield 25% degree of substitution (DS25)^32,33^. First, UP VLVG alginate was dissolved in 2-(N-morpholino)ethanesulfonic acid (MES, Thermo Scientific, P15J509)) buffer at 1% w/v (pH 6.5, 0.1M MES, 0.3M UP NaCl) for 60 minutes stirring at 400RPM. Second, N-hydroxysuccinimide (NHS, Sigma Aldrich)) and 1-ethyl-3-(3-dimethylaminopropyl)-carbodiimide hydrochloride (EDC, Sigma Aldrich), were immediately dissolved at 1/20 the reaction volume in MES buffer and added dropwise in 2.5X molar excess based on the number of carboxylic acid groups on each alginate monomer to the reaction vessel. After 15 minutes (to enable sufficient activation of carboxylic acid groups by EDC/NHS), Tz-HCl (dissolved and sonicated for 60 minutes at 1/20 the reaction volume in MES buffer), was added dropwise to the reaction vessel at 1mmol per gram of alginate, stirred at 400RPM overnight at room temperature for 18 hours, and protected from light with aluminum foil.

The reaction product was then collected in falcon tubes at a volume of ≤ 35mL and ultracentrifuged at 10,000RPM for 60 minutes to pellet unreacted Tz-HCl and crashed reaction side products. After ultracentrifugation, the product was decanted, filtered (0.22μm filter), and purified via dialysis utilizing a 3.5 kDa MW cutoff membrane (VWR International, 20020178) using a decreasing salt gradient from 0.15M to 0.10M then 0.05M to 0M UP NaCl (VWR Life Sciences, 24B2656648) in 4L of milliQ water, with a salt-solution exchange every 2 hours with one overnight step for 4 days. Dialyzed DS25 Tz-VLVG alginate was then treated with activated carbon (Thermo Scientific, P31I026) at a 1:1 alginate:carbon mass ratio for 30 minutes stirring at 400RPM and then sterile filtered (0.22μm filter), and lyophilized for long-term storage at -20ºC, protected from light.

### Primary human CD34^+^ hematopoietic stem cell isolation from cord blood and cell culture expansion

Primary CD34^+^ human hematopoietic stem cells (HSCs) were isolated from cord blood units procured from the Dana Farber Cancer Institute Pasquarello Tissue bank. HSCs were isolated via density gradient centrifugation with Lymphoprep and SepMate-50 tubes (StemCell Technologies), followed by positive isolation (EasySep™ Human Cord Blood CD34 Positive Selection Kit III, StemCell Technologies), and frozen in freezing media (BAMBANKER™) at -80ºC overnight prior to long-term storage in liquid nitrogen at a density of 2e5-2e6 cells/mL. Prior to hydrogel encapsulation for 3D culture on day 0, cord blood-derived HSCs were expanded for 7 days on 6 well suspension culture plates (GenClone, 25-100) at standard tissue culture conditions with a seeding density of 1e4 cells/well in an expansion media comprised of IMDM, GlutaMAX™ (Gibco) supplemented with 20% BIT 9500 (StemCell Technologies), 1% penicillin-streptomycin (Thermo), carrier free recombinant human (rh) SCF (100 ng/mL, R&D), rh Flt3L (100 ng/mL, R&D), and rh TPO (50 ng/mL, R&D), 1 μg/mL human low-density lipoproteins (hLDL, Calbiochem, 437644), 24μM beta-mercaptoethanol, 60μM L-ascorbic acid (Sigma Aldrich, A4544-25G), and 5mM CaCl_2_.

### Trans-cyclooctene modification of DLL-4 and VCAM-1 ligands

TCO-PEG_5_-DLL-4 and TCO-PEG_5_-VCAM-1 were prepared via maleimide-thiol bioconjugation chemistry, tethering TCO to the Fc-tag of either rh Fc-DLL-4 (Sino Biological, 10171-H02H), or rh Fc-VCAM-1 (Sino Biological, 10113-H02H). Briefly, rh Fc-DLL-4 or Fc-VCAM-1 was diluted in PBS to 0.5 mg/mL (6.173μM and 4.921 μM, respectively), and reduced with 209μM of Tris(2-carboxyethyl)phosphine hydrochloride (TCEP HCl, Sigma Aldrich, 51805-45-9) for 20 minutes while rocking on a nutating mixer at room temperature, followed by the addition of 418μM of Trans-cyclooctene-polyethylene-glycol_5_-maleimide (TCO-PEG_5_-mal, ConjuProbe, CP-6009) at a molar ratio of 1:34 of Fc-tag DLL-4 or VCAM-1:TCEP HCl and 1:2 molar ratio of TCEP HCl:TCO-PEG_5_-mal. The bioconjugation reaction was then incubated overnight at 4ºC on a nutating mixer prior to purification in IMDM, GlutaMAX^™^ (containing 1% penicillin-streptomycin) via a protein concentrator column (Pierce, MWCO 30kDa, 88502) as per the manufacturer’s protocol. Purified stock concentrations of TCO-PEG_5_-DLL-4 or TCO-PEG_5_-VCAM-1 were then measured via nanodrop to quantify protein concentration and A280, prior to short-term storage at 4ºC or long-term storage at -80ºC.

### DLL-4 and VCAM-1 conjugation and CD34^+^ HSC encapsulation in thymus-mimetic extracellular matrices of tunable viscoelastic properties

TCO-PEG_5_-DLL-4 and TCO-PEG_5_-VCAM-1 were reacted with DS25 Tz-VLVG alginate in a solution of phenol red HyClone™ Hank’s 1X Balanced Salt Solution (HBSS, Cytiva, containing 20mM HEPES) in sterile 1.5mL Eppendorf tubes at a mass ratio of 20 μg:4 μg:1 mg TCO-PEG_5_-DLL-4:TCO-PEG_5_-VCAM-1:Tz-VLVG alginate overnight at 4ºC on a nutating mixer prior to IPN hydrogel-based ETN fabrication and CD34^+^ HSC encapsulation for 3D cell culture as previously described with some modifications^32,33^. Briefly, rat tail telo-collagen type I (8 mg/mL in 25mM acetic acid, Corning) was neutralized on ice at a stock neutralized concentration of 6 mg/mL to 6.5 < pH < 7.0 and added to the protein-polymer conjugate solution chilled on ice. Calcium carbonate slurry (CaCO_3_) was then obtained by high-speed vortexing precipitated CaCO_3_ nanoparticles (nano-PCC, Multifex-MM, Specialty Minerals) at a stock concentration of 100 mg/mL in sterile water (Water-For-Injection, WFI, Cytiva) and added to the pre-gel solution for a final concentration of: 2 mg/mL collagen I, 1.25% w/v DLL-4/VCAM-1 modified Tz-VLVG alginate, and 10mM or 100mM CaCO_3_ (soft and stiff, respectively). CD34^+^ HSCs were retrieved from culture and suspended to 52.5e6 cells/mL in HBSS (containing 20mM HEPES) on ice and added to the pre-gel solution and gently vortexed for a final suspension concentration of 3.5e6 cells/mL.

Immediately before gelation, an appropriate amount (4x molar excess with respect to the CaCO_3_ concentration) of freshly dissolved glucono-delta-lactone (GDL, EMD Millipore, 0.4 mg/μL in HBSS (20mM HEPES)) was added to the pre-gel solution and gently vortexed to induce gelation. To form elastic gels, 400μM of multivalent 4-arm-PEG_5k_-Norbornene (4a-PEG_5k_-Nb, Creative PEGWorks, stock concentration of 25 mg/mL in HBSS (20mM HEPES)) was then immediately added to the pre-gel solution and gently vortexed to reinforce the matrix with covalent crosslinks. The resulting solutions were then immediately cast into a 48-multiwell MatTek dish (MatTek Corporation, P48G-1.5-6-F) by micropipette in 50μL volumes. Gelation was then allowed at 37ºC in the tissue culture incubator for 60 minutes, followed by equilibration in 500μL of HBSS (20mM HEPES, 5mM CaCl_2_) at 37ºC for an additional 60 minutes, prior to replacement of buffer with 400μL of human progenitor T-cell differentiation media.

3D cell culture of CD34^+^ HSCs in staged human progenitor T-cell differentiation media

After 3D ETN encapsulation, 3D cell culture was carried out in a serum free and fully defined differentiation media comprised of IMDM, GlutaMAX™ (Gibco) supplemented with 20% BIT 9500 (StemCell Technologies), 1% penicillin-streptomycin (Thermo), carrier free rh SCF (24 ng/mL, R&D), rh Flt3L (9 ng/mL, R&D), rh IL-3 (5 ng/mL, R&D), rh IL-7 (10 ng/mL, R&D), rh TNF-α (5 ng/mL, R&D), rh CXCL12 (10 ng/mL, R&D), 1 μg/mL human low-density lipoproteins (hLDL, Calbiochem, 437644), 24μM beta-mercaptoethanol, 60μM L-ascorbic acid (Sigma Aldrich, A4544-25G), and 5mM CaCl_2_ on day 0 – day 7 (stage I media).

For 3D cell culture on day 7 – day 14 (stage II), 3D cell culture was carried out in IMDM, GlutaMAX™ (Gibco) supplemented with 20% BIT 9500 (StemCell Technologies), 1% penicillin-streptomycin (Thermo), carrier free rh SCF (120 ng/mL, R&D), rh Flt3L (8 ng/mL, R&D), rh IL-3 (1 ng/mL, R&D), rh IL-7 (45 ng/mL, R&D), rh TNF-α (0.4 ng/mL, R&D), rh CXCL12 (15 ng/mL, R&D), 1 μg/mL human low-density lipoproteins (hLDL, Calbiochem, 437644), 24μM beta-mercaptoethanol, 60μM L-ascorbic acid (Sigma Aldrich, A4544-25G), and 5mM CaCl_2_.

### Rheological characterization of thymus-mimetic extracellular matrices

Rheological characterization of the IPN hydrogel-based ETNs was performed on a stress-controlled rheometer (AR-G2, TA instruments) using a 20mm diameter 1º tilt angle cone geometry as previously described^32,33,60^. IPN-hydrogel solution (90μL) was immediately loaded onto the Peltier plate after pulse vortexing. A solution of HBSS containing 20mM HEPES and 5mM CaCl_2_ was used to wet the periphery of the cone geometry along with a humidity chamber to prevent dehydration. Oscillatory rheology was used to measure the storage modulus (G’), loss modulus (G’’), and loss angle (O) after equilibrium was achieved. Rheological tests included an oscillatory time sweep for 3 hours at 25ºC (1 Hz, 1% strain), followed by an oscillatory frequency sweep (1% strain, 0.1-10 Hz). A shear stress relaxation test was then performed with a 15% shear strain and strain rise time of 1 second, and then maintaining a fixed 15% shear strain over time for 10 hours. Creep/plasticity measurements were performed by first performing a 1 hour oscillatory time sweep at 37ºC (1 Hz, 1% strain), and then by applying an instantaneous 100 Pa fixed shear stress for 1 hour, followed by an instantaneous 0 Pa fixed shear stress for 2 hours at 20ºC.

### Cell retrieval from thymus-mimetic extracellular matrices

Hydrogels were first washed in 500μL of PBS, and subsequently digested with 200μL of 100 U/mL collagenase IV (Worthington Biochemical Corp., NC9919937) and 25 U/mL alginate lyase (Sigma Aldrich, A1603) in IMDM, GlutaMAX^™^ (containing 1% penicillin-streptomycin), at 37ºC for 15 minutes, followed by trituration with a P1000 pipette and an additional 10 minutes of incubation at 37 ºC. Thereafter, additional trituration with a P1000 pipette was performed to fully dissolve gels. The collagenase and lyase were then quenched with 200μL of IMDM, GlutaMAX^™^ (containing 20% BIT 9500, 1% penicillin-streptomycin, 1 μg/mL hLDL, 24μM beta-mercaptoethanol, 60μM L-ascorbic acid and 5mM CaCl_2_), after which cells were collected and then passed through 40μm mini-strainers (pluriSelect USA, 43-10040-40) into sterile 1.5mL Eppendorf tubes. Collected cells were pelleted (at 1600RPM for 3 minutes), washed in PBS twice, and then resuspended for downstream analysis (e.g., flow cytometry or total RNA isolation for RT-qPCR).

### Flow cytometry analysis

Cells isolated from thymus-mimetic matrices were transferred to 96-well v-bottom plates and kept at 4ºC for immunostaining. Cells were stained with LIVE/DEAD™ fixable blue cell stain (invitrogen, L34961) according to the manufacturer’s protocol. Cells were then subsequently blocked with human FcX Fc receptor blocking solution (BioLegend, 422301) for 15 minutes and stained with surface protein-binding antibodies for 30 minutes. eBioscience™ Flow cytometry staining buffer (invitrogen, 00-4222-26) was used during staining. Flow cytometry was then performed with a BD FACSymphony^™^ A3 flow cytometer. Gating was performed based on fluorescence-minus-one controls. The complete set of antibodies used for flow cytometry is listed in **Supplementary Table 2** in the supporting information.

### Small molecule inhibition studies

Small molecule inhibitors targeting features of the notch1 pathway and actomyosin contraction were purchased from Selleckchem. Inhibitor working concentrations were determined using the CellTiter-Blue® cell viability assay (Promega, G8080) with CD34^+^ HSCs cultured on 96-well U-bottom suspension plates in expansion media containing 0.5% DMSO for 7 days, with a media exchange on day 4. Determined working concentrations and catalog numbers for the small molecule inhibitors are provided in **Supplementary Table 3** in the supporting information.

### RNA isolation and RT-qPCR gene expression analysis

CD34^+^ HSCs were cultured in thymus-mimetic matrices for 96 hours in stage I media. Cells were then retrieved from matrices for subsequent total RNA isolation. Total RNA was isolated via PureLink™ RNA Micro Scale Kit (Invitrogen, 12183016A) according to the manufacturer’s protocol. Purity and concentration of isolated total RNA was assessed via nanodrop, prior to long-term storage at -80ºC. cDNA was then synthesized from total RNA using SuperScript III Reverse Transcriptase (Invitrogen, 18080093), according to the manufacturer’s protocol, and stored at -80ºC. cDNA was amplified with respective primers utilizing FastStart Essential SYBR Green Master Mix (Roche, 06402712001). Thermocycling and quantification was performed on a CFX-96 RT-PCR thermocycler (Bio-rad) as according to the manufacturer’s protocol (Roche) with two technical replicates for each biological replicate. Relative expression of individual genes was calculated by the delta cycle and delta-delta cycle threshold (2^-ΔCt^and 2^-ΔΔCt^) methods with the expression of GAPDH as internal reference housekeeping genes. PCR primer sequences are available in **Supplementary Table 4** in the supporting information.

### Statistical analyses

Statistical analyses were performed in Prism Graphpad v.10.2.3. Statistical tests used one-way ANOVA and post-hoc tests for multiple comparisons, two-way ANOVA and post-hoc tests for multiple comparisons, or unpaired *t* test with Welch’s correction for comparison between two groups. P-values < 0.05 were considered significant unless otherwise noted. Error bars represent the standard deviation of the mean unless otherwise noted.

## Conclusions

Mechanically confining IPN hydrogel-based 3D cell culture comprised of collagen type I and alginate polysaccharides functionalized with DLL-4 and VCAM-1 was used as a model 3D ETN to manipulate human progenitor (pro)T-cell differentiation through orthogonal changes in modulus and stress relaxation behavior. Soft-viscous 3D ETN matrices permitted activation of the T-cell development GRN, and enhanced the emergence of T-competent progenitor cells from encapsulated CD34^+^ HSCs through synergies between notch1-signaling and actomyosin contractility via integrin-substrate adhesions. Stiffer, elastic matrices inhibit HSC commitment to the T-lineage and proT-cell proliferation, and rather promoted the generation of Myeloid-cells. These observations emphasize potential mechanical determinants of signaling pathways indispensable to thymopoiesis, and highlights ECM mechanics as a variable in controlling hematopoietic stem cell fate decisions within their cellular niche. These findings may have utility for future *in vitro* models of thymopoiesis and therapeutic T-cell biomanufacturing strategies from stem cells.

## Acknowledgements

We thank Dr. Bryan Nerger (Harvard University), Dr. Yuesong Hu (Wyss Institute), Dr. Grace Bingham (Harvard University), Dr. Yoav Binenbaum (Wyss Institute), Mason Dacus (Harvard University), Dr. Asel Primbetova (University of British Columbia), Dr. Peter W. Zandstra (University of British Columbia), Dr. Cedric Tremblay (University of Manitoba), and Dr. Yale S. Michaels (University of Manitoba) for insightful scientific discussions. We thank Michael Carr (Wyss Institute) and Alexander Pauer (Wyss Institute) for technical staff assistance. We thank Dr. Matthew Kerr (Lightning Biotherapeutics) and Siddharth Iyer (Wyss Institute) for assistance with the RT-qPCR experiments. We thank the Dana Farber Cancer Institute Pasquarello tissue bank for procuring and providing umbilical cord blood units used for experimentation. We are especially grateful to the patients who donated their umbilical cord-blood samples used in these studies. We acknowledge funding from the NIH NCI (Wyss Institute i3 center, Wellcome Leap Foundation (HOPE), and NSF (GRFP, Harvard MRSEC). W.-H.J. was supported by the NIH NCI (K00CA253759). N.L-G. was supported by the CRIS Cancer Foundation.

