## Supplementary Information for "Human progenitor T-cell differentiation regulated by the mechanical resistance of thymus-mimetic extracellular matrices"

| **Condition** | **Tz-VLVG** (% w/v) | **Collagen I** (mg/mL) | **CaCO_3_** (mM) | **GDL** (mM) | **TCO-PEG_5_-DLL-4** (μg/mL) | **TCO-PEG_5_-VCAM-1** (μg/mL) | **4-a-PEG5k-Nb** (μM) | **Storage Modulus** (kPa) | **Loss angle** (º) |
| --- | --- | --- | --- | --- | --- | --- | --- | --- | --- |
| Soft-viscous | 1.25 | 2 | 10 | 40 | 250 | 50 | 0 | 0.40 | 6.0 |
| Soft-elastic | 1.25 | 2 | 10 | 40 | 250 | 50 | 400 | 0.40 | 4.5 |
| Stiff-viscous | 1.25 | 2 | 100 | 400 | 250 | 50 | 0 | 1.00 | 6.0 |
| Stiff-elastic | 1.25 | 2 | 100 | 400 | 250 | 50 | 400 | 1.00 | 4.5 |

**Supplementary Table 1. Physicochemical compositions of thymus-mimetic extracellular matrices**

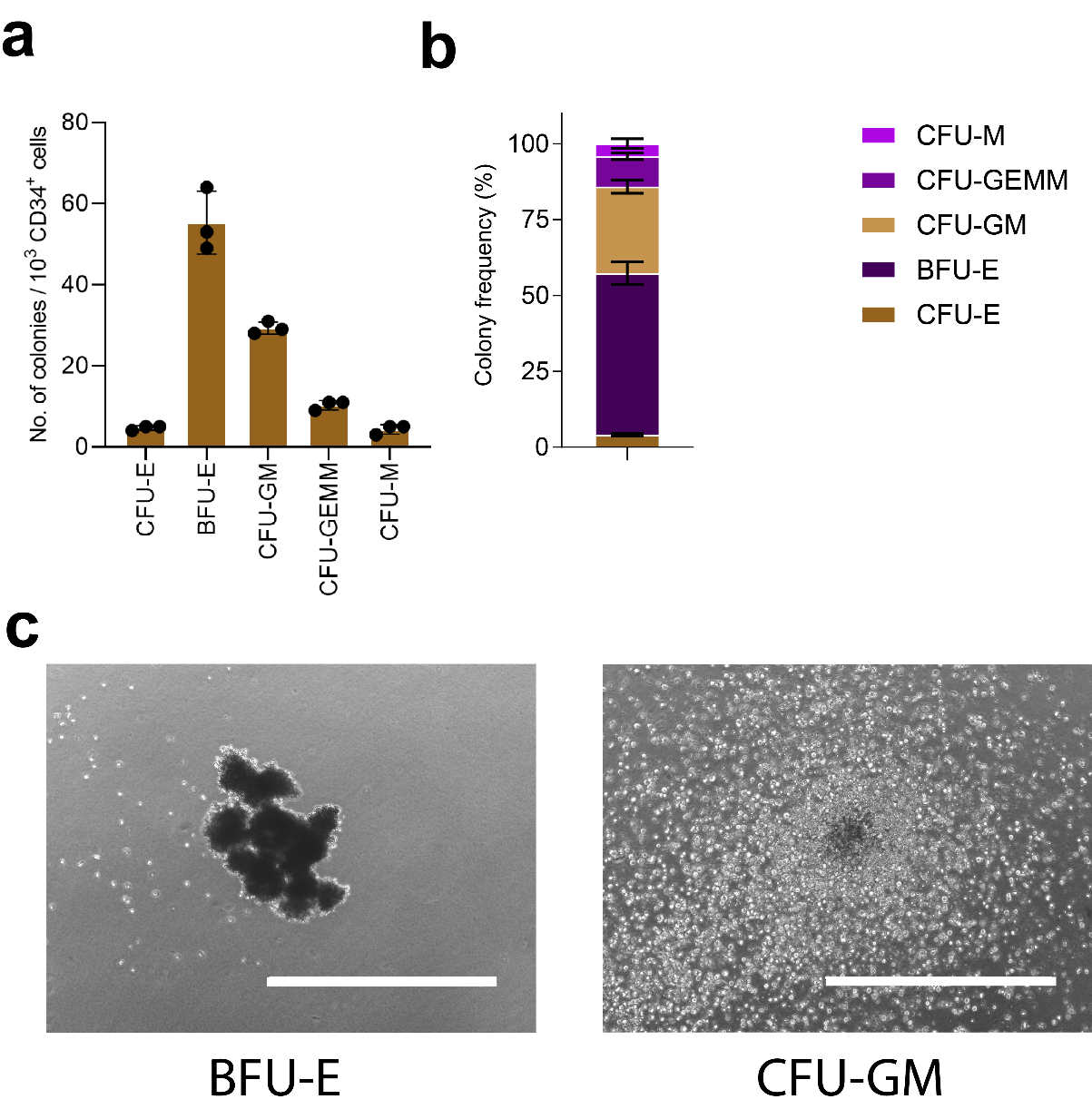

**Supplementary Fig 1. Colony forming potential of cord blood-isolated CD34^+^ HSCs cultured in** **MethoCult™ SF H4636.** a) Number of colony forming units (CFUs) per 10^3^ input CD34^+^ cells (n = 3). b) Frequency of colonies in CFU assay (n = 3). c) Representative brightfield images of BFU-E and CFU-GM in CFU assay after 14 days of incubation. CFU-E: Colony forming unit-erythrocyte, BFU-E: Burst forming unit-erythrocyte, CFU-GM: Colony forming unit-granulocyte-macrophage, CFU-GEMM: Colony forming unit-granulocyte-erythrocyte-monocyte-megakaryocyte, CFU-M: Colony gorming unit-macrophage. Data is visualized as mean values +/- SD. Scale bar: 1000μm.

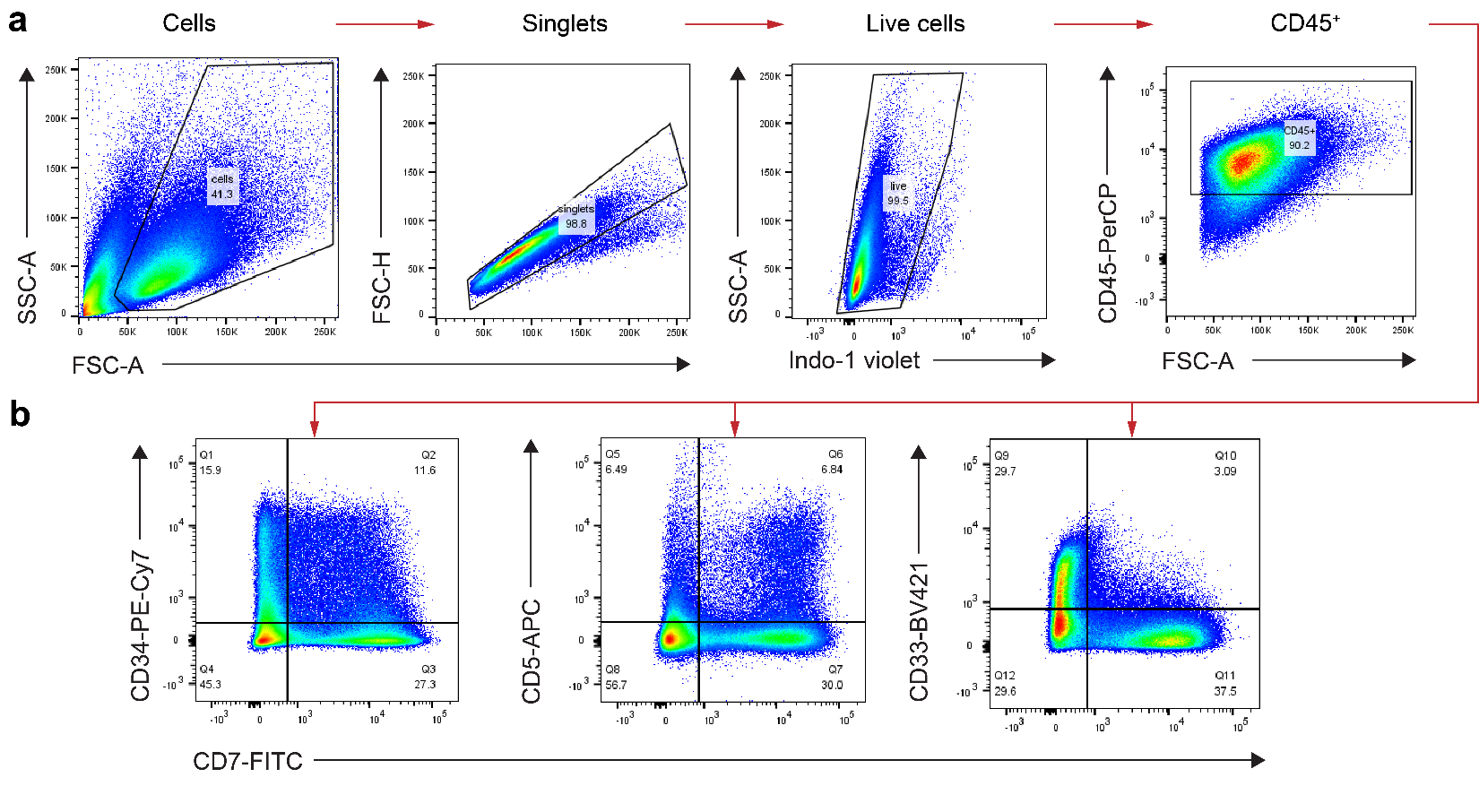

**Supplementary Fig. 2. Flow cytometry gating strategy.** a) Gating of live CD45^+^ cells. b) Gating of CD34^+/-^, CD7^+/-^, CD5^+/-^ and CD33^+/-^ progenitor cells on live CD45^+^ cells.

| **Antigen** | **Fluorophore** | **Isotype - Clone** | **Distributor** | **Catalog no.** |
| --- | --- | --- | --- | --- |
| CD45 | PerCP | Mouse - 2D1 | Biolegend | 368506 |
| CD34 | PE-Cy7 | Mouse - 580 | Biolegend | 343516 |
| CD7 | FITC | Mouse - 6B7 | Biolegend | 343104 |
| CD5 | APC | Mouse - UCHT2 | Biolegend | 300612 |
| CD33 | BV421 | Mouse – P67.6 | Biolegend | 366622 |
| Live/Dead | Indo-1 Violet | N/A | invitrogen | L34961 |

**Supplementary Table 2. List of antibodies and fluorescent dyes used for flow cytometry studies.**

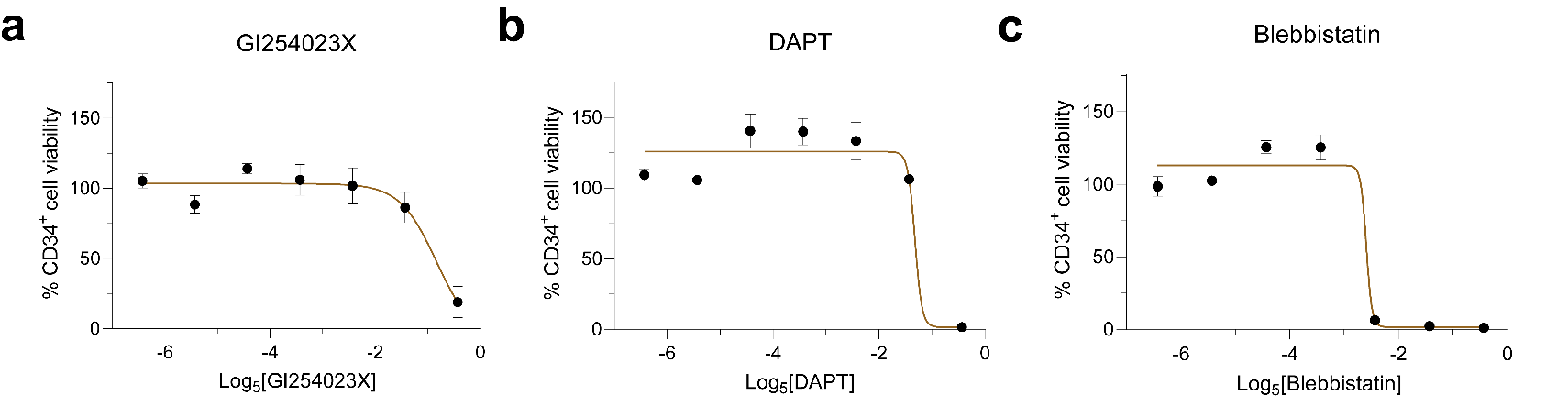

**Supplementary Fig. 3. Determination of pharmacological inhibitor working concentrations for small molecule inhibition of notch1 signaling and actomyosin contraction.** a) CD34^+^ HSC viability as a function of GI254023X concentration (n = 3). b) CD34^+^ HSC viability as a function of DAPT concentration (n = 3). c) CD34^+^ HSC viability as a function of Blebbistatin concentration (n = 3). Working concentrations were screened from 0.032μM-500μM of inhibitor across seven 5-fold serial dilutions. Data is visualized as mean values +/- SD.

| **Inhibitor** | **Target** | **Stock concentration** (in DMSO) | **Working concentration** | **Distributor** | **Catalog no.** |
| --- | --- | --- | --- | --- | --- |
| GI254023X | ADAM10 | 100mM | 20μM | Selleckchem | S8660 |
| DAPT | γ-secretase | 100mM | 20μM | Selleckchem | S2215 |
| Blebbistatin | myosin II | 100mM | 4μM | Selleckchem | E1249 |

**Supplementary Table 3. Small molecule inhibitors for pharmacology studies.**

| **Gene** | **Species** | **Forward primer** | **Reverse primer** |
| --- | --- | --- | --- |
| Deltex | Human | ATCGGAGAAGGCTCTACAGG | CGTCTGGCCTCCTTTCTAACT |
| TCF7 | Human | TGCACATGCAGCTATACCCAG | TGGTGGATTCTTGGTGCTTTTC |
| Notch1 | Human | GAGGCGTGGCAGACTATGC | CTTGTACTCCGTCAGCGTGA |
| Hes1 | Human | CCTGTCATCCCCGTCTACAC | CACATGGAGTCCGCCGTAA |
| Pu.1 | Human | TGCAATGTCAAGGGAGGGGG | AAACCCTTCCATTTTGCACGC |
| E2a | Human | CCGACTCCTACAGTGGGCTA | CGCTGACGTGTTCTCCTCG |
| GAPDH | Human | GTCTCCTCTGACTTCAACAGCG | ACCACCCTGTTGCTGTAGCCAA |

**Supplementary Table 4. Forward and reverse primers for RT-qPCR analysis.**

| **Cytokine** | **Expansion media**  (d-7-d0) | **Stage I media**  (d0-d7) | **Stage II media** (d7-d14) | **Distributor** | **Catalog No.** |
| --- | --- | --- | --- | --- | --- |
| SCF | 100 ng/mL | 24 ng/mL | 120 ng/mL | R&D Systems | 7466SC010CF |
| Flt3L | 100 ng/mL | 9 ng/mL | 8 ng/mL | R&D Systems | 308FK010CF |
| TPO | 50 ng/mL | - | - | R&D Systems | 288TP005CF |
| IL-3 | - | 5 ng/mL | 1 ng/mL | R&D Systems | 203IL050CF |
| IL-7 | - | 10 ng/mL | 45 ng/mL | R&D Systems | 207IL010CF |
| TNF-α | - | 5 ng/mL | 0.4 ng/mL | R&D Systems | 103707588CF |
| CXCL12 | - | 10 ng/mL | 15 ng/mL | R&D Systems | 103714126 |

**Supplementary Table 5.** **Cytokine compositions of CD34^+^ HSC expansion media, stage I proT-cell differentiation media, and stage II proT-cell differentiation media.**

**Supplementary materials and methods**

Colony forming unit assay

Cryopreserved HSCs were thawed and resuspended in IMDM, GlutaMAX™ (Gibco) supplemented with 20% BIT 9500 (StemCell Technologies), 1% penicillin-streptomycin (Thermo), 1 μg/mL human low-density lipoproteins (hLDL, Calbiochem, 437644), 24μM beta-mercaptoethanol, and 60μM L-ascorbic acid (Sigma Aldrich, A4544-25G) at a density of 1e4 cells/mL prior to culture in MethoCult™ SF H4636 (StemCell Technologies) as according to the manufacturer’s protocol with some modifications. Briefly, resuspended cells were diluted 1:10 in 2.0mL sterile Eppendorf tubes containing 1.5mL of MethoCult™ and vortexed vigorously for 5 seconds to evenly disperse cells and achieve a cell density of 1e3 cells/mL. The suspension was allowed to stand on ice for 5 minutes to allow bubbles to rise. 1.1mL of suspension was subsequently drawn into a primed sterile 3mL luer lock syringe attached to a 16G-needle and dispensed onto a 35mm culture dish (StemCell Technologies, 27150). After even coating of suspension, 35mm culture dishes were placed into a sterile 100mm petri dish along with one open 35mm culture dish containing 3mL of sterile DI H_2_O to prevent dehydration. The MethoCult™ cultures were then incubated at standard tissue culture conditions for 14 days. After incubation, colonies were counted manually on an inverted brightfield microscope under 4X objective with a STEMgrid™-6 plate (StemCell Technologies, 27000).
